# Low antigen abundance limits efficient T-cell recognition of highly conserved regions of SARS-CoV-2

**DOI:** 10.1101/2021.10.13.464181

**Authors:** Srividhya Swaminathan, Katie E. Lineburg, George R. Ambalathingal, Pauline Crooks, Emma J. Grant, Sonali V Mohan, Jyothy Raju, Archana Panikkar, Laetitia Le Texier, Zheng Wei Marcus Tong, Keng Yih Chew, Michelle A. Neller, Kirsty R. Short, Harsha Gowda, Stephanie Gras, Rajiv Khanna, Corey Smith

## Abstract

Understanding the immune response to severe acute respiratory syndrome coronavirus (SARS-CoV-2) is critical to overcome the current coronavirus disease (COVID-19) pandemic. Efforts are being made to understand the potential cross-protective immunity of memory T cells, induced by prior encounters with seasonal coronaviruses, in providing protection against severe COVID-19. In this study we assessed T-cell responses directed against highly conserved regions of SARS-CoV-2. Epitope mapping revealed 16 CD8^+^ T-cell epitopes across the nucleocapsid (N), spike (S) and ORF3a proteins of SARS-CoV-2 and five CD8^+^ T-cell epitopes encoded within the highly conserved regions of the ORF1ab polyprotein of SARS-CoV-2. Comparative sequence analysis showed high conservation of SARS-CoV-2 ORF1ab T-cell epitopes in seasonal coronaviruses. Paradoxically, the immune responses directed against the conserved ORF1ab epitopes were infrequent and subdominant in both convalescent and unexposed participants. This subdominant immune response was consistent with a low abundance of ORF1ab encoded proteins in SARS-CoV-2 infected cells. Overall, these observations suggest that while cross-reactive CD8^+^ T cells likely exist in unexposed individuals, they are not common and therefore are unlikely to play a significant role in providing broad pre-existing immunity in the community.

## Introduction

Severe acute respiratory syndrome coronavirus 2 (SARS-CoV-2) is responsible for the current coronavirus disease 2019 (COVID-19) global pandemic. The virus has infected more than 190 million individuals worldwide, causing more than 4 million death as of July 2021 [1]. SARS-CoV-2 belongs to the beta-coronavirus family. The four most common human coronaviruses include the beta coronaviruses HCoV-OC43 and HCoV-HKU-1, and alpha coronaviruses HCoV-229E and HCoV-NL63. Infection with seasonal coronaviruses is typically associated with mild symptoms. In comparison, SARS-CoV-1, SARS-CoV-2 and MERS-CoV display a range of disease symptoms ranging from asymptomatic to life threating conditions [2, 3]. It remains unclear whether prior exposure to common circulating coronaviruses provides individuals with pre-existing immunity that is capable of protecting them against COVID-19.

The genomic structure of SARS-CoV-2 has several open reading frames (ORF) responsible for virus entry and replication inside the host cells. ORF1ab is a polyprotein responsible for viral RNA replication and transcription. Additional ORFs encode the structural proteins which include the nucleocapsid (N), spike (S), envelope (E) and membrane (M) and six accessory proteins (ORF3a, ORF6, ORF7a, ORF7b, ORF8 and ORF10) [4, 5]. Recent studies have defined the T-cell immunodominance hierarchies of these antigens [6, 7]. Both CD4^+^ and CD8^+^ T cell responses are induced against most proteins, with immunodominant CD8^+^ T-cell responses targeting ORF3a, N and S, and CD4^+^ T-cell responses primarily directed against N, S and M. Immunodominant T-cell epitopes have also been defined in other proteins, including ORF1ab [8-12]. The induction of both antigen-specific CD4^+^ and CD8^+^ T cells have been associated with protection from severe COVID-19 [13]. In spite of the protective immunity resulting from either SARS-CoV-2 infection or vaccination, the ongoing emergence of SARS-CoV-2 variants will likely result in new variants that are capable of evading these immune responses. A number of studies have suggested that the existence of cross-protective memory T-cell responses, induced following prior encounters with seasonal coronaviruses, may play some protective role against severe COVID-19 [10, 14-19]. We recently demonstrated cross-reactive immunity towards a single highly conserved immunodominant CD8^+^ T-cell epitope [14, 20]. Meanwhile most other studies have focused on cross-recognition between CD4^+^ T-cell epitopes that typically display less conservation in amino acid sequence [2, 9]. In this study we sort to assess whether highly conserved regions within coronaviruses harbor CD8^+^ T-cell epitopes that are capable of being cross-recognized between different coronaviruses and may provide cross-protection against SARS-COV-2 by way of pre-existing T-cell immunity. We demonstrate that the ORF1ab protein does contain highly conserved T-cell epitopes that can be recognised by T cells from both COVID-19 convalescent and unexposed participants. However, the responses to these epitopes are not present in the majority of human leukocyte antigen (HLA) matched participants and are typically subdominant when detectable. More importantly, we show that this lack of consistent T-cell reactivity against highly conserved regions is coincident with low abundance of ORF1ab-encoded proteins in virus-infected cells.

## Results

### Conservation of predicted T-cell epitopes in beta coronaviruses

In order to test whether pre-existing memory CD8^+^ T cells, generated from prior exposure to circulating beta-coronaviruses, were capable of recognizing homologous SARS-CoV-2 epitopes we set out to identify highly conserved regions within SARS-CoV-2 proteins. Here, we aligned the genome sequences of HCoV-OC43 and SARS-CoV-2 and identified sequences that displayed 100% homology and were a minimum of 8 amino acids in length. While we identified only one conserved region between ORF3a, ORF6, ORF7a, ORF7b, ORF8 and ORF10 which is known to contain the HLA-B^*^07:02 restricted epitope SPRWYFYYL encoded in the nucleocapsid protein, we noted that ORF1ab contains 27 of these highly conserved regions (Figure. 1; indicated as red vertical lines). These highly conserved sequences within ORF1ab were encoded from nsp4 onwards; and the majority were found in nsp12-16. To assess the potential immunogenicity of these regions we designed 43 overlapping peptides of 12-13 amino acids in length that covered the entire sequence within these regions (Table 1). The comparison of SARS-Cov2 T-cell epitopes to the corresponding HCoV-OC43 sequences demonstrated a maximum of 2 amino acid changes (Table 1). Computational HLA class I epitope predictions for 18 common alleles revealed 163 potential strong binding (Score - <1%) HLA class I-restricted epitopes encoded within the conserved 12-mer to 13-mer sequences (Supplementary Table 1).

**Figure 1.**
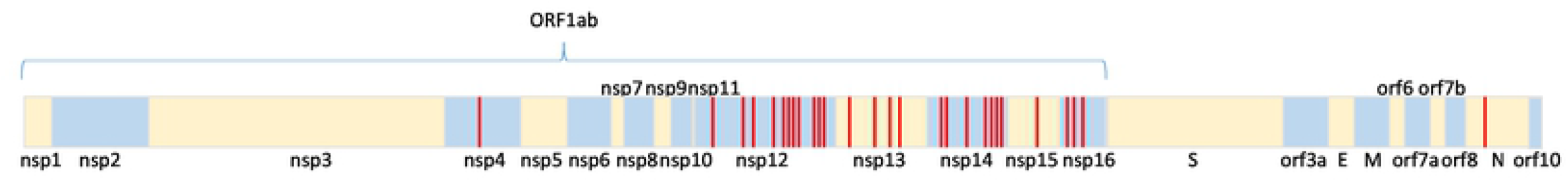
Sequence conservation between SARS-CoV-2 and HCoV-OC43. A schematic representation of the conserved regions of SARS-CoV-2 that are of a minimum length for recognition by CD8+ T cells. Regions with a minimum sequence of eight conserved amino acids between SARS-CoV-2 and HCoV-OC43 are highlighted in red.

**Table 1:**
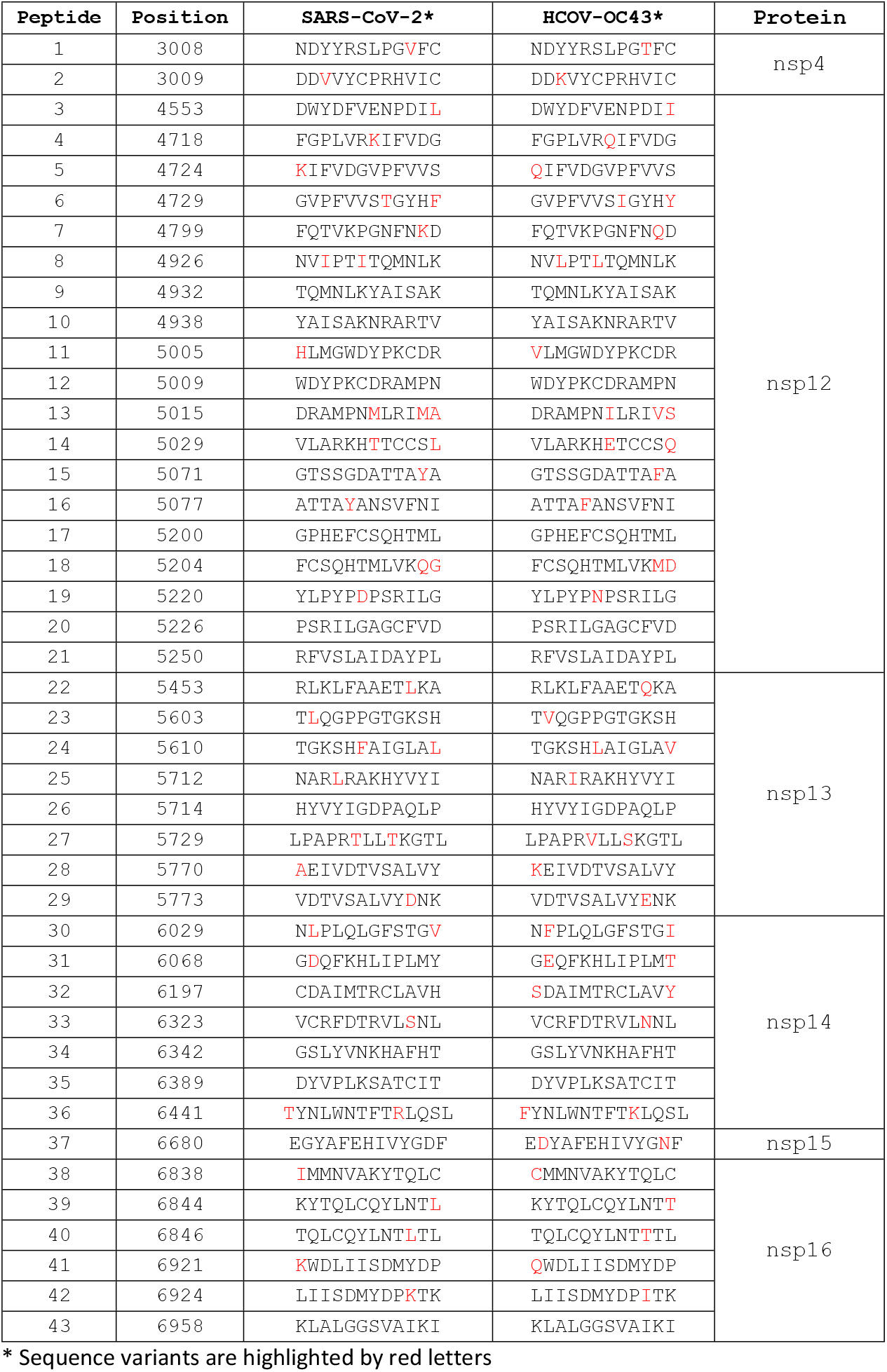
Peptides encoded in the conserved orf1ab peptide pool.

| Peptide | Position | SARS-CoV-2* | HCOV-OC43* | Protein |
| --- | --- | --- | --- | --- |
| 1 | 3008 | NDYYRSLPGVFC | NDYYRSLPGTFC | nsp4 |
| 2 | 3009 | DDVVYCPRHVIC | DDKVYCPRHVIC |  |
| 3 | 4553 | DWYDFVENPDIL | DWYDFVENPDI I | nsp12 |
| 4 | 4718 | FGPLVRKIFVDG | FGPLVRQIFVDG |  |
| 5 | 4724 | KIFVDGVPFVVS | QIFVDGVPFVVS |  |
| 6 | 4729 | GVPFVVS TGYHF | GVPFVVS IGYHY |  |
| 7 | 4799 | FQTVKPGNFNKD | FQTVKPGNFNQD |  |
| 8 | 4926 | NV IPTITQMNLK | NV LPTLTQMNLK |  |
| 9 | 4932 | TQMNLKYAISAK | TQMNLKYAISAK |  |
| 10 | 4938 | YAISAKNRARTV | YAISAKNRARTV |  |
| 11 | 5005 | HLMGWDYPKCDR | VLMGWDYPKCDR |  |
| 12 | 5009 | WDYPKCDRAMPN | WDYPKCDRAMPN |  |
| 13 | 5015 | DRAMPNMLRIMA | DRAMPN ILRIVS |  |
| 14 | 5029 | VLARKH TTCCSL | VLARKH ETCCSQ |  |
| 15 | 5071 | GTSSGDATTAYA | GTSSGDATTAF A |  |
| 16 | 5077 | ATTAYANSVFNI | ATTAF ANSVFNI |  |
| 17 | 5200 | GPHEFCSQHTML | GPHEFCSQHTML |  |
| 18 | 5204 | FCSQHTMLVKQG | FCSQHTMLVKMD |  |
| 19 | 5220 | YLPYPDPSRILG | YLPYPNPSRILG |  |
| 20 | 5226 | PSRILGAGCFVD | PSRILGAGCFVD |  |
| 21 | 5250 | RFVSLAIDAYPL | RFVSLAIDAYPL |  |
| 22 | 5453 | RLKLFAAETLKA | RLKLFAAETQKA | nsp13 |
| 23 | 5603 | TLQGPPGTGKSH | TVQGPPGTGKSH |  |
| 24 | 5610 | TGKSHFAIGLAL | TGKSHLAIGLAV |  |
| 25 | 5712 | NARLR AKHYVYI | NAR I RAKHYVYI |  |
| 26 | 5714 | HYVYIGDPAQLP | HYVYIGDPAQLP |  |
| 27 | 5729 | LPAPR TLLTKGTL | LPAPR VLLSKGTL |  |
| 28 | 5770 | AEIVDTV SALVY | KEIVDTV SALVY |  |
| 29 | 5773 | VDTV SALVYDNK | VDTV SALVYENK |  |
| 30 | 6029 | NLPLQLGFSTGV | NFPLQLGFSTGI | nsp14 |
| 31 | 6068 | GDQFKHLIPLMY | GEQFKHLIPLMT |  |
| 32 | 6197 | CDAIMTRCLAVH | SDAIMTRCLAVY |  |
| 33 | 6323 | VCRFDTRVL SNL | VCRFDTRVLNNL |  |
| 34 | 6342 | GSLYVNKHAFHT | GSLYVNKHAFHT |  |
| 35 | 6389 | DYVPLKSATCIT | DYVPLKSATCIT |  |
| 36 | 6441 | TYNLWNTFTRLQSL | FYNLWNTFTKLQSL |  |
| 37 | 6680 | EGYAFEHIVYGDF | EDYAFEHIVYGNF | nsp15 |
| 38 | 6838 | IMMNVAKYTQLC | CMMNVAKYTQLC | nsp16 |
| 39 | 6844 | KYTQLCQYLNTL | KYTQLCQYLNTT |  |
| 40 | 6846 | TQLCQYLNTLTL | TQLCQYLNTTTL |  |
| 41 | 6921 | KWDLIISDMYDP | QWDLIISDMYDP |  |
| 42 | 6924 | LIISDMYDPKTK | LIISDMYDPITK |  |
| 43 | 6958 | KLALGGSVAIKI | KLALGGSVAIKI |  |

\* Sequence variants are highlighted by red letters

### CD8^+^ T-cell recognition of SARS-CoV-2 antigens

To assess the potential immunogenicity of these peptides, we combined all 43 conserved peptides into a single peptide pool (ORF1ab-C), and used this to stimulate PBMC from both SARS-CoV-2 exposed and unexposed individuals. PBMC were cultured for 14 days in the presence of interleukin-2 and, following recall with the same ORF1ab-C pool, were assessed for the presence of IFN-γ producing CD8^+^ T cells using ICS and flow cytometric analysis (Supplementary Figure 1). Participants were also assessed for the presence of T-cell responses directed against overlapping peptide pools (OPPs) from ORF3a, N and S. Due to the limited detection of responses toward ORF1ab-C peptides, we also assessed T-cell responses directed against nsp12 in 51 SARS-CoV-2 convalescent volunteers and 8 unexposed volunteers. Immunodominant T-cell responses in convalescent individuals were predominantly detected against ORF3a, N and S, with limited recognition of ORF1ab-C and nsp12 (Figure. 2A). The majority of convalescent individuals generated a CD8^+^ T-cell IFN-γ response above a threshold of 0.5% towards ORF3a (n=42/60), N (n=45/60), and Spike S1 (n=43/60), with 41% reacting to S2 (n=25/60) (Figure 2B&C). In contrast, only 15% of convalescent participants showed an ORF1ab-C specific CD8^+^ IFN-γ response >0.5% (n=9/60), and 37% showed an nsp12-specific response (19/51). Although we only screened a small number of unexposed participants, our observations suggested a different pattern of antigen recognition in these donors, with very few displaying CD8^+^ T-cell responses above the 0.5% threshold in response to ORF3a (n=0/8), N (n=1/8), S1 (n=1/8) and S2 (n=1/8); and more frequent recognition of ORF1ab-C (n=3/16) and nsp12 (n=4/8) (Figure 2D&E).

**Figure 2.**
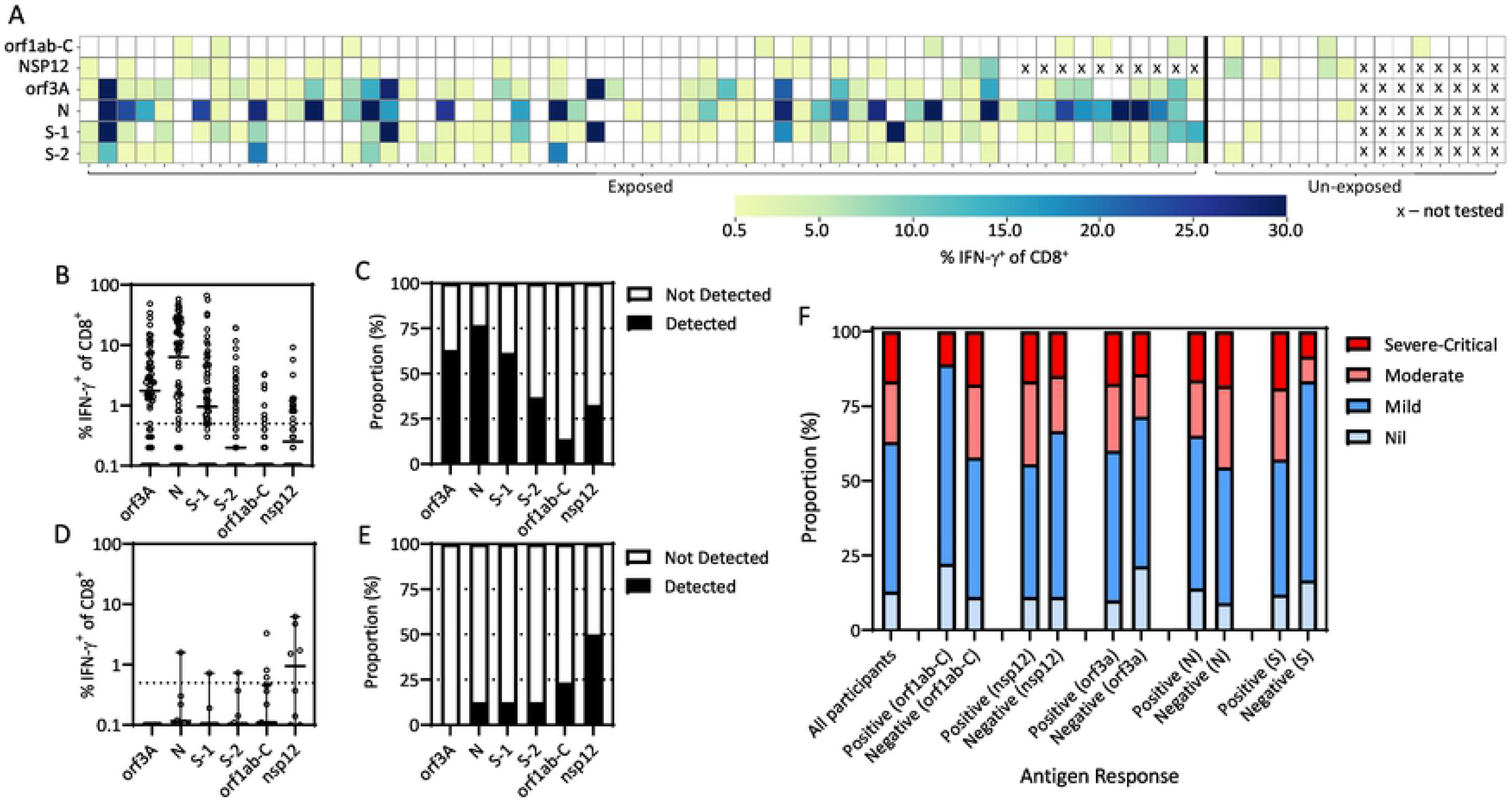
CD8+ T cell response to conserved regions of ORF1ab in COVID-19 convalescent participants. **(A)** PBMC from convalescent and unexposed participants were stimulated with a pool of peptides covering the ORF1ab conserved regions (see Table 1) and overlapping peptide pools of SARS-CoV-2 (nsp12, orf3a, N, S1 and S2) were cultured for two weeks in the presence of IL-2, then assessed for IFN-γ production following recall with the peptide pool. Data represents CD8^+^ IFN-γ^+^ producing cells to each peptide pools from convalescent participants ORF1ab-C, ORF3a, N, S-1, S-2 (n=60); nsp12 (n=51) and unexposed participants ORF1ab-C (n=16); nsp12, ORF3a, N, S-1, S-2 (n=8). The participants not tested with the peptide pools is denoted as “X” in the Heatmap. The responses greater than 0.5% of CD8^+^ IFN-γ^+^ producing cells to the peptide pools were considered positive. **(B)** The dot plot represents frequency of CD8^+^ IFN-γ^+^producing cells in convalescent participants responding to the overlapping peptide pools (ORF3a, N, S-1, S-2 ORF1ab-C and NSP-12). Responses were considered positive when the number of IFN-γ^+^ CD8^+^ T cells was greater than 0.5%. **(C)** The proportional analysis were carried out in convalescent participants to the overlapping peptide pools (ORF3a, N, S-1, S-2 ORF1ab-C and nsp12) are shown in the bar graph. Data represents proportion of responders to non-responders across overlapping peptide pools (ORF3a, N, S-1, S-2 ORF1ab-C and nsp12) **(D)** The percentage of CD8^+^ IFN-γ^+^ producing cells in unexposed participants (n=18) responding to the overlapping peptide pools (ORF3a, N, S-1, S-2 ORF1ab-C and nsp12) are shown in the dot plot. Responses were considered positive when the number of IFN-γ^+^ CD8^+^ T cells was greater than 0.5%. **(E)** The proportional analysis were carried out in unexposed participants to the overlapping peptide pools (ORF3a, N, S-1, S-2 ORF1ab-C and nsp12) are shown in the bar graph. The number of responses detected and not detected to each overlapping peptide pools (ORF3a, N, S-1, S-2 ORF1ab-C and nsp12) is represented in the bar graph. **(F)** The histogram represents the comparison of CD8^+^ IFN-γ^+^ producing cells as positive response (> 0.5%) and negative response (< 0.5%) across SARS-CoV-2 antigens (ORF1ab-C, nsp12, ORF3a, N, S). The antigen responses were compared to the symptoms (nil, mild, moderate and severe to critical) from the convalescent participants.

We next stratified our convalescent participants based upon their COVID-19 severity (nil, mild, moderate or severe/critical) and assessed the proportion responding to each of the SARS-CoV-2 antigens (ORF1ab-C, nsp12, ORF3a, N, S) (Figure. 2F). Among the 60 convalescent volunteers, seven had no symptoms, 27 had mild symptoms, 11 had moderate symptoms and nine had severe-critical symptoms. Disease information was not available for the remaining seven participants. Relative to all participants analysed, an increased proportion of those who generated an ORF1ab-C response had mild or no symptoms (89% ORF1ab-C responders to 63% non-responders). Conversely, volunteers lacking a spike protein response demonstrated a higher incidence of mild or no symptoms (83%) compared to those with spike responses (63%). No other potential associations were evident between participants’ symptoms and response to the other antigens. In addition, we saw no association between the presence of an ORF1ab-C or nsp12 CD8^+^ T-cell response and the magnitude of the neutralising antibody response (Supplementary Figure 2).

### Epitope mapping reveals CD8^+^ T-cell epitopes from SARS-CoV-2 antigens

To define the minimal epitope determinates in each of the peptide pools we generated a custom peptide matrix to identify responding peptides from the ORF1ab-C pool, and using a standard matrix for the nsp12, ORF3a, N and S, [14, 21]. Following minimization and HLA-restriction assays we were able to identify five CD8^+^ T-cell epitopes encoded within the ORF1ab-C and nsp12 (Figure 3A). These five epitopes included the previously reported <u>SARS</u>-CoV-1 and SARS-CoV-2 epitopes <u>FVD</u>GVPFVV (HLA-A^*^02:07-restricted), <u>LPY</u>PDPSRI (B^*^51:01-restricted) and DTDFVNEFY (A^*^01:01-restricted) and two novel epitopes, YAFEHIVY (HLA-B35:01-restricted) and NVIPTITQMNL (A^*^02:05-restricted). We also identified 16 CD8^+^immunodominant T-cell epitopes from the ORF3a, N and S proteins (Supplementary Table 2). Representative flow cytometry analysis is presented for five of these CD8^+^ T-cell epitopes, <u>TPS</u>GTWLTY (B^*^35:01-restricted), <u>SPR</u>WYFYYL (B^*^07:02-restricted-restricted) and <u>MEV</u>TPSGTWL (B^*^40:01-restricted) from N, <u>FTS</u>DYYQLY (A^*^01:01-restricted) from ORF3a, and <u>YLQ</u>PRTFLL (A^*^02:01-restricted) from S [14, 22, 23] (Figure 3B).

**Figure 3.**
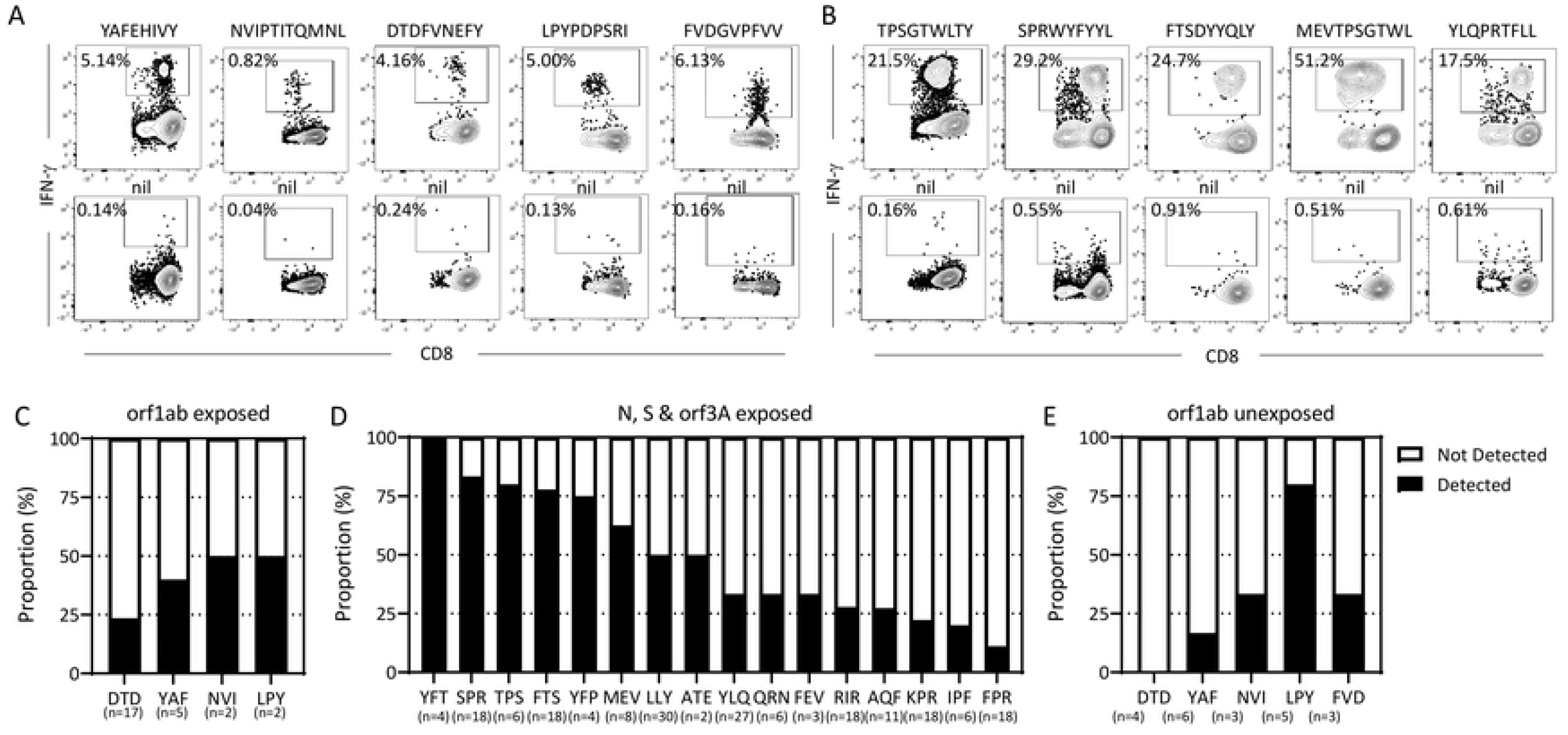
Identification of CD8+ T cells epitopes in the conserved regions of ORF1ab. Minimal peptide epitope sequences from each antigen were determined using a standard peptide matrix, followed by peptide minimization and HLA restriction.. **(A)** Representative flow cytometric plots of IFN-γ production by CD8^+^ T cells following recognition of peptides from ORF1ab (YAF, NVI, DTD, LPY, FVD). The flow cytometric plots of each CD8+ T cell epitope were compared to nil (without peptide) to demonstrate epitope-specific IFN-γ production **(B)** Representative flow cytometric plots of IFN-γ production by CD8^+^ T cells following recognition of peptides from N (TPS,SPR, MEV); S-1 (YLQ); ORF3a (FTS). **(C)** Data represents the proportion of responders and to non-responders to each of ORF1ab epitopes (DTD, YAF, NVI, LPY) from convalescent participants, where (n) is the number of HLA-matched participants tested for each epitope. **(D)** Data represents the percentage proportion of responders and non-responders to each of the epitopes from N (SPR, TPS, MEV, ATE, QRN, RIR, AQF, KPR, FPR); S (YFP, YLQ, FEY, IPF) & ORF3a (YFT, FTS, LLY) from convalescent participants, where (n) is the number of HLA-matched participants tested for each epitope. **(E)** Data represents the percentage proportion of responders to non-responders to each of the epitopes from ORF1ab (DTD, YAF, NVI, LPY, FVD) from unexposed participants, where (n) is the number of HLA-matched participants tested for each epitope.

**Table 2:** Conservation of ORF1ab SARS-CoV-2 peptides by variant.

|  |  | Variants of Concern |  |  |  | Variants of Interest |  |  |  |  |  |
| --- | --- | --- | --- | --- | --- | --- | --- | --- | --- | --- | --- |
| Position | Sequence | Alpha | Beta | Gamma | Delta | Epsilon | Zeta | Eta | Theta | Iota | Kappa |
| 6682-6689 | Y<u>A</u>FEHIV<u>Y</u> | 99.92 | 100.00 | 100.00 | 99.08 | 99.92 | 100.00 | 100.00 | 100.00 | 99.91 | 100.00 |
| 5221-5229 | LPYPDPSR<u>I</u> | 99.93 | 92.74 | 99.98 | 100.00 | 99.90 | 99.78 | 100.00 | 100.00 | 99.91 | 100.00 |
| 4926-4936 | NV<u>I</u>PTITQM<u>N</u>L | 99.93 | 100.00 | 100.00 | 100.00 | 100.00 | 100.00 | 100.00 | 100.00 | 99.97 | 100.00 |
| 4726-4734 | FVDGVPFV<u>V</u> | 99.96 | 100.00 | 99.98 | 100.00 | 99.85 | 100.00 | 100.00 | 100.00 | 100.00 | 100.00 |
| 5130-5138 | DTDFVNEF<u>Y</u> | 99.69 | 100.00 | 99.49 | 100.00 | 99.62 | 100.00 | 100.00 | 100.00 | 99.74 | 100.00 |
| 1637-1646 | TTDPSFLGR<u>Y</u> | 99.17 | 100.00 | 99.66 | 23.55 | 99.68 | 100.00 | 100.00 | 100.00 | 99.73 | 99.11 |
| 3886-3894 | KLWAQCVQ<u>L</u> | 99.95 | 100.00 | 99.98 | 100.00 | 99.97 | 99.78 | 98.97 | 100.00 | 99.99 | 100.00 |

To determine the proportion of convalescent volunteers that respond to the ORF1ab-C epitopes, we stimulated available HLA-matched PBMC with the cognate peptide for 14 days and assessed IFN-γ production. Participants were considered responders if >0.5% of CD8^+^ T cells produced IFN-γ following subtraction of the background (no peptide) control. Consistent with our peptide pool responses, a lower proportion of HLA-matched participants responded to the ORF1ab-C/nsp12 encoded epitopes (Figure 3C) compared to the immunodominant epitopes defined in ORF3a, N or S (Figure 3D). To perform a similar assessment in unexposed individuals we cultured PBMC from unexposed donors with HLA-matched ORF1ab encoded epitopes and expanded for 14 days in the presence of IL-2. T cells were then recalled with cognate peptide and assessed for IFN-γ production. We were able to detect T-cell responses against four of the five ORF1ab epitopes in unexposed participants (Figure 3E), excluding the HLA-A^*^01:01 restricted DTD epitope which was mapped from the nsp12 peptide pool rather than the ORF1ab-C pool. While a majority of B^*^51:01 positive volunteers (4/5) responded to the LPY epitope, we only detected responses to the HLA B^*^35:01-restricted YAF, HLA A^*^02:05-restricted NVI and HLA A^*^02:07-restricted FVD epitopes in one individual for each, further confirming the rarity of T-cell responses to these epitopes in individuals exposed to either SARS-CoV-2 or common circulating coronaviruses.

### Conservation of SARS-CoV-2 encoded epitopes in circulating coronaviruses

Despite the low prevalence of T cells that recognize ORF1ab epitopes in both COVID-19 convalescent and unexposed individuals, our observations suggest that the high degree of conservation within this ORF1ab region, between SARS-CoV-2 and common circulating coronaviruses, is associated with cross-recognition. Comparative sequence analysis with circulating coronaviruses demonstrated a high level of conservation in the four epitopes (YAF, LPY, NVI, FVD) that unexposed volunteers responded to (Figure 4, Supplementary Table 3). In contrast, the DTD epitope, which was not recognised by unexposed volunteers, together with the epitopes mapped in other regions of SARS-CoV-2, displayed less homology with circulating coronaviruses. In addition, two previously published immunodominant epitopes <u>TTD</u>PSFLGRY (HLA-A^*^01:01-restricted) and <u>KL</u>WAQCVQL (HLA-A^*^02:01-restricted), which are encoded in the nsp3 and nsp7 proteins of ORF1ab, also display limited conservation in circulating coronaviruses, further emphasizing the rare nature of conserved SARS-CoV-2 encoded CD8^+^ T-cell epitopes in circulating coronaviruses [24, 25]. To explore conservation within the ORF1ab encoded epitopes of SARS-CoV-2 variants, we assessed sequence variation in 10,000 recent isolates from each of the 10 different variants (Table 2). Similar to the high level of conservation observed between different circulating coronaviruses, most of the epitopes displayed a high level of conservation in SARS-CoV-2 isolates, with the majority displaying >99% sequence homology in all isolates. More frequent changes were evident in two epitopes: In the HLA-B^*^51:01-restricted LPY epitope, 7% of SARS-CoV-2 alpha isolates contained an L to F substitution in position (P) 1; and in the HLA-A^*^01:01-restricted TTD epitope, 76% of delta isolates contained a P to I substitution at P4 (Supplementary Table 4).

**Figure 4.**
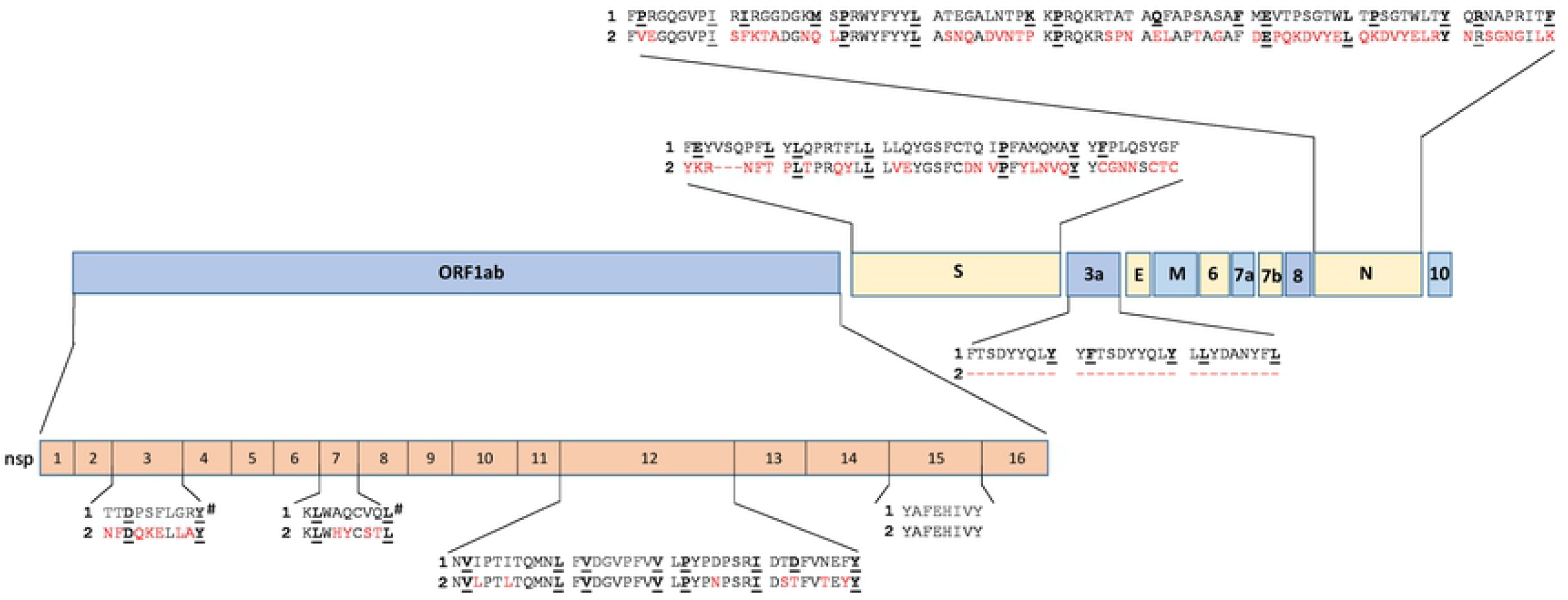
Alignment of SARS-CoV-2 peptide epitopes with their corresponding peptide sequence in HCov-OC43. The peptide epitope sequences are aligned with their location in the genomic structure of SARS-CoV-2. Differences in the peptide sequences between SARS-CoV-2 (**1**) and HCoV-OC43 (**2**) are highlighted in red and HLA anchoring positions are underlined. # Represent two previously published ORF1ab epitopes not tested in this study

While the ORF1ab-C epitopes displayed a higher level of conservation, there was still a number of amino acid differences between SARS-CoV-2 and circulating coronaviruses that could impact cross-recognition. To assess the impact of these amino acid changes on T-cell activation, we stimulated LPY, YAF, NVI and DTD-specific T cells generated from both convalescent and unexposed volunteers with variant peptides and assessed IFN-γ production. As expected, due to our inability to generate T cells from SARS-CoV-2 unexposed HLA-A^*^01:01^+^ individuals, DTD-specific T cells did not recognise any of the variant epitopes present in other circulating coronaviruses (Figure 5A). LPY-specific T cells from both exposed and convalescent volunteers did not efficiently cross-react with the OC43-encoded variant despite a single D to N amino acid substitution at P5 (Figure 5B). Similarly, YAF-specific T cells were unable to cross-react with the HKU1 variant despite a conserved E to D substitution at P4, however YAF-specific T-cell cultures did show some reactivity towards the NL63 and 229E variant, FNFEHVVY, which encodes three amino acid substitutions, including an A to N change at the P2 anchor residue (Figure 5C). The NVI-specific T cells generated from a single unexposed donor displayed reactivity against all variants assessed (Figure 5D). To further assess the potential cross-reactivity of YAF and NVI specific T cells, we used a peptide titration of each variant to assess T-cell activation. Despite recognition of the NL63 and 229E variant FNFEHVVY at high peptide concentrations, YAF-specific T cells showed markedly reduced functional avidity against this variant peptide sequence (Figure 5E), suggesting that this cross-reactivity is unlikely to be physiologically relevant. Similarly, NVI-specific T cells displayed low functional avidity against the variant peptides (Figure 5F). However, we did note the high functional avidity of the YAF and NVI-specific T cells against their cognate peptide. These observations further emphasise the fine-specificity that CD8^+^ T cells have for their cognate peptide and demonstrate the complexity of peptide cross-recognition against substitutions at different positions in the peptide sequence.

**Figure 5.**
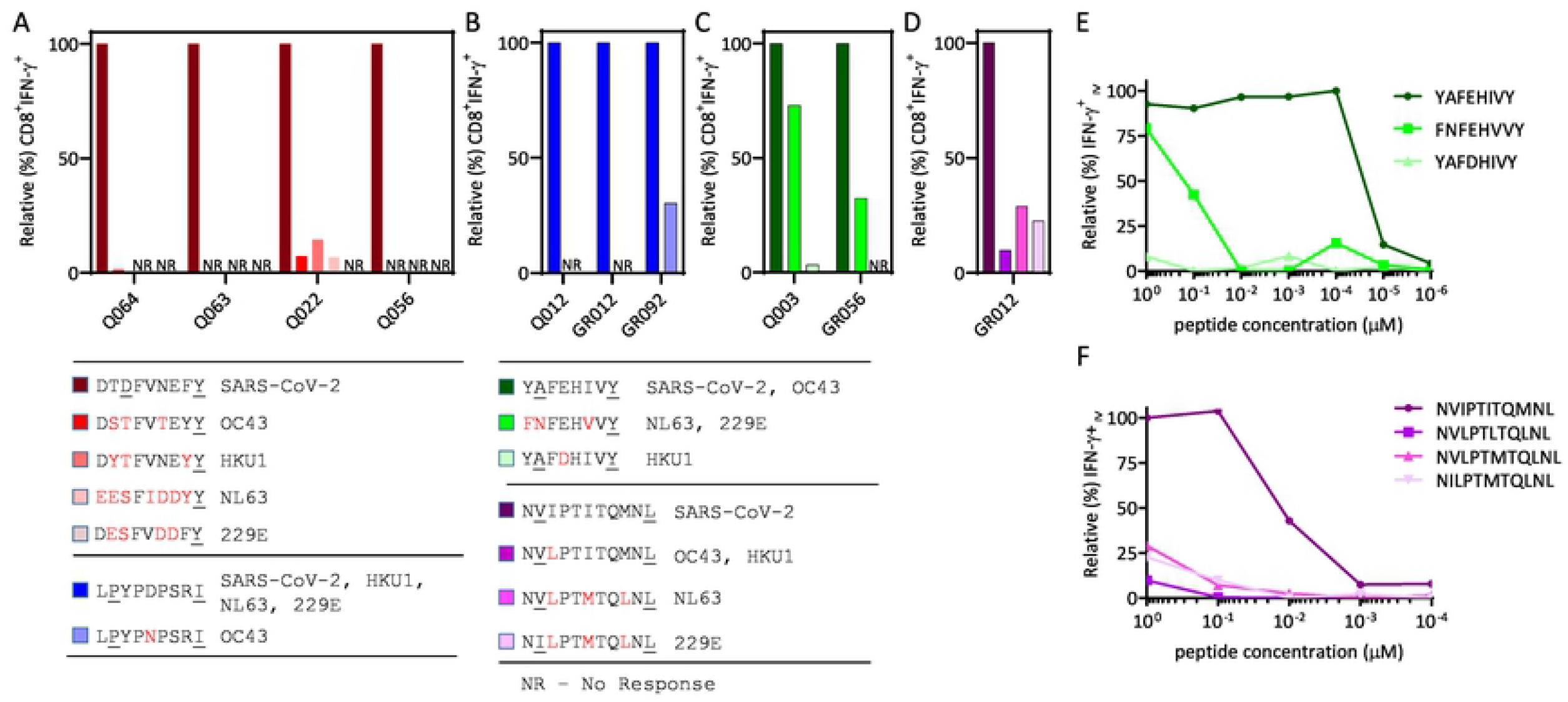
CD8+ T cell responses and functional avidity of ORF1ab peptides derived from pandemic and seasonal coronaviruses. T cells specific for ORF1ab encoded epitopes (YAF, NVI, LPY, DTD) were assessed for their ability to recognise variant peptide sequences found in common circulating coronaviruses using an ICS assay. **(A)** CD8^+^IFN-γ responses of LPY-specific T cells from COVID-19 exposed (Q012) and unexposed participants (GR012, GR092) following recall stimulation with LPYPDPSRI (SARS-CoV-2, HKU-1, NL63, 229E) and LPYPNPSRI (OC43). **(B)** CD8^+^ IFN-γ responses of YAF-specific T cells from COVID-19 exposed (Q003) and unexposed participants (GR056) following recall stimulation with YAFEHIVY (SARS-CoV-2, OC43), FNFEHVVY (NL63, 229E) and YAFDHIVY (HKU-1). **(C)** CD8^+^IFN-γ responses of DTD-specific T cells expanded from COVID-19 exposed participants (Q022, Q056,Q063, Q064) following recall stimulation with DTDFVNEFY (SARS-CoV-2), DSTFVTEYY (OC43), DYTFVNEYY (HKU-1), EESFIDDYY (NL63) and DESFVDDFY (229E) peptides in an ICS assy. **(D)** CD8 IFN-γ responses of NVI-specific T cells expanded from an unexposed donor (GR012) following recall stimulation with NVI PTITQMNL (SARS-CoV-2), NVLPTLTQMNL (OC43, HKU-1), NVLPTMTQLNL (NL63) and NILPTMTQLNL (229E). **(E)** T cells from Q003 were stimulated with ten-fold serial dilutions of the YAF peptide variants and assessed for IFN-γ production using ICS. Data is shown as the relative percentage of IFN-γ producing CD8+ T cells compared to maximal stimulation.**(F)** T cells from GR012 were stimulated with ten-fold serial dilutions of the NVI peptide variants and assessed for IFN-γproduction using ICS. Data is shown as the relative percentage of IFN-γ producing CD8+ T cells compared to maximal stimulation.

### Antigen abundance limits recognition of conserved ORF1ab epitopes to T cells with high avidity

While low functional avidity of T-cell clones selected against the ORF1ab epitopes may explain the subdominant nature of the T-cell responses against these epitopes, our preliminary assessment of T-cell avidity in YAF and NVI specific T cells suggest that this is unlikely. To confirm this we assessed functional avidity in ORF1ab-specific T cells and compared this to T cells specific for the immunodominant TPS, SPR, MEV, FTS and YLQ epitopes from ORF3a, N and S. We then calculated the concentration of peptide required to induce 50% maximal activation of CD8^+^ T cells (EC50). Analysis of the EC50 from both convalescent and unexposed T cells demonstrated a consistent high sensitivity to ORF1ab-C peptide, with a median of 0.0012µM (range: 3×10e-5 to 0.14) (Figure. 6A). The EC50 from ORF3a, N and S had a median of 0.013µM (range: 0.0002 to 0.5971) (Figure 6B), indicating that the immunodominance observed in these epitopes was not simply driven by selection of T-cell clones with higher functional avidity.

**Figure 6.**
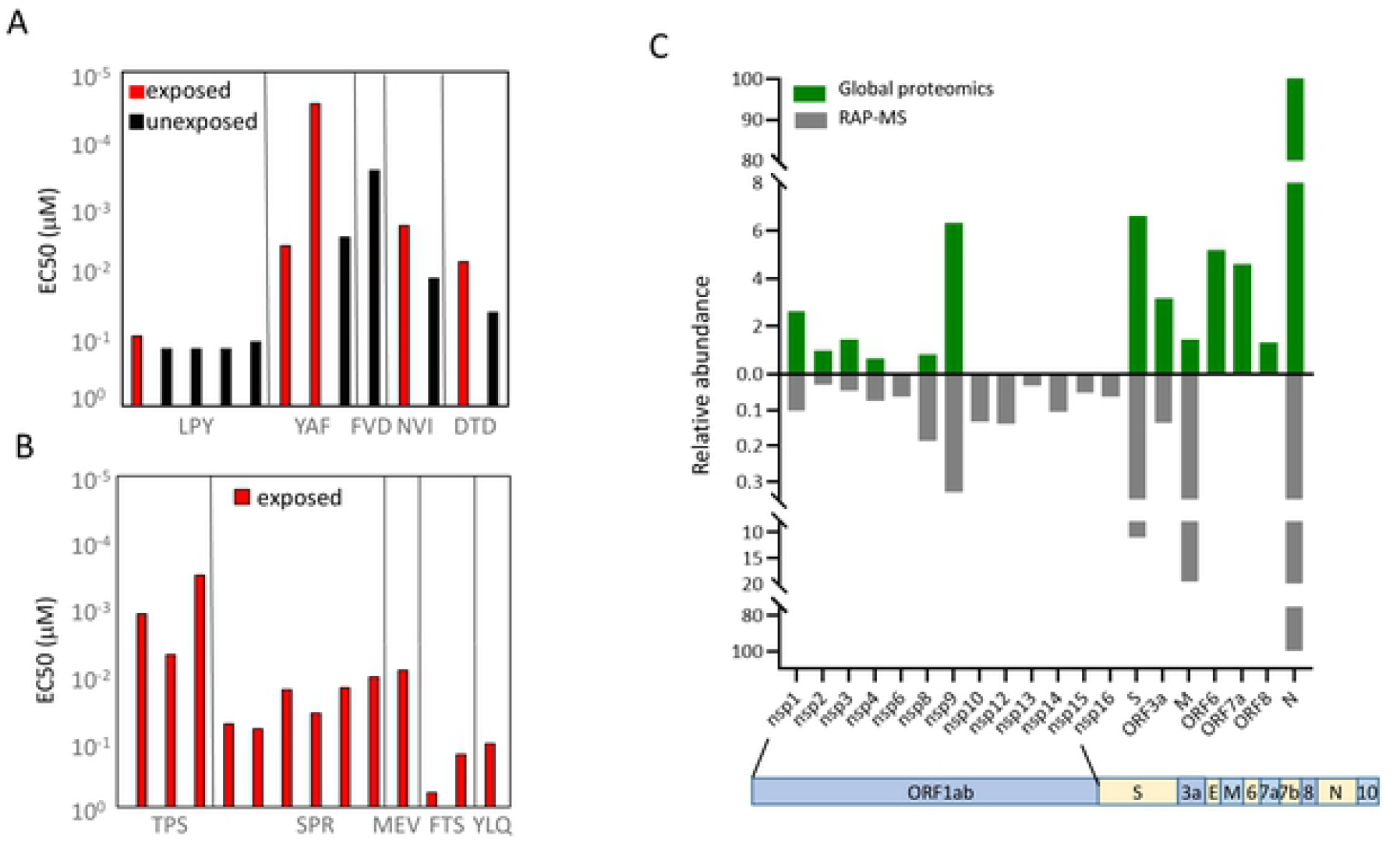
Effective concentration and antigen abundance across SARS-CoV-2 antigens. **(A)** Data represents the avidity at the effective concentration used to induce half maximal response (EC50) of ORF1ab-C, nsp12 peptides from convalescent participants (LPY n=1, YAF n=2, NVI n=1, DTD n=1) and unexposed participants (LPY n=4, YAF n=1, FVD n=1, NVI n=1, DTD n=1). **(B)** Data represents the avidity at the effective concentration used to induce half maximal response (EC50) of N, S-1, ORF3a peptides from convalescent participants (TPS n=3, SPR n=6, MEV n=1, FTS n=2, YLQ n=1). Mann-Whitney test was used to perform statistical analysis. **(C)** Relative abundance of SARS-CoV-2 proteins in infected cells. Histogram at the top shows relative abundance of SARS-CoV-2 proteins in infected cells using global proteomics analysis. Histogram at the bottom shows relative abundance of SARS-CoV-2 proteins in infected cells following enrichment of RNA-binding proteins using RNA antisense purification coupled with mass spectrometry (RAP-liMS).

We have previously reported that the efficiency of recognition of endogenous antigen can impact the immunodominance hierarchy of virus-specific T cells in humans [26]. Although we were not able to explore endogenous presentation of the CD8^+^ T-cell epitopes by SARS-CoV-2 infected T cells, due to an absence of appropriate target cells, we examined abundance of virus encoded proteins in SARS-CoV-2 infected cells from three independent studies [27-30]. In all three studies assessed, the nsp10-16 proteins of ORF1ab could not be detected in global proteomic analysis (Figure 6C, Supplementary Figure 3). Nsp10-16 were only detectable following enrichment of RNA-binding proteins. In contrast all three studies demonstrated high abundance of the ORF3a, N and S proteins [27-30].

Overall, our observations demonstrate that although highly conserved regions of coronaviruses do provide a source of epitopes for T-cell cross-recognition, limited antigen abundance in infected cells likely restricts T-cell priming in coronavirus exposed individuals, thus limiting widespread protection in the population by pre-existing CD8^+^ T cells.

## Discussion

Previous observations have suggested that exposure to seasonal coronaviruses may provide a level of protection against COVID-19 through the activation of T cells capable of cross-recognizing highly conserved epitopes found in both SARS-CoV-2 and seasonal coronaviruses [10, 14-19]. Recent observations from our group supported this contention by demonstrating the presence of cross-reactive T-cell immunity in COVID-19 convalescent and unexposed individuals against a single highly conserved immunodominant HLA B^*^07:02-restricited CD8^+^ T-cell epitope encoded in the N protein of SARS-CoV-2 [14]. In the current study we extended these observations to explore other highly conserved regions of SARS-CoV-2 encoded within the large ORF1ab polyprotein. However, unlike the immunodominant nature of the responses directed against the conserved HLA B^*^07:02-restricted SPR epitope, immune responses directed against the conserved ORF1ab epitopes were infrequent and typically subdominant in frequency. These observations suggest that while cross-reactive CD8^+^ T cells likely exist in unexposed individuals, they are not common and therefore are unlikely to play a significant role in providing broad pre-existing immunity in the community.

While rare in our cohort, T cells that could cross-recognize the conserved regions of ORF1ab did appear to be more prevalent in participants who were asymptomatic or had mild symptoms. Two recent studies support a potential protective role for these T cells against COVID-19. Mallajosyula et al. demonstrated that patients with milder COVID-19 had CD8^+^ T cells specific for conserved epitopes [31]. Swadling et al. demonstrated that abortive infection, which does not lead to seroconversion, is associated with the expansion of pre-existing T cells [32]. In both instances the conserved CD8^+^ T-cell epitopes were predominantly encoded in ORF1ab. Earlier studies by Grifoni et al. also demonstrated that ORF1ab and nsp12 in particular provided a hotspot for CD8^+^ T-cell responses in unexposed individuals [6]. These observations suggest that the conserved regions of ORF1ab across all coronaviruses could offer targets for broad vaccine based immunity.

Despite the potential role in limiting disease severity it was clear that the conserved regions of ORF1ab induced uncommon subdominant T-cell responses. This is consistent with previous studies assessing T-cell responses against the whole genome of SARS-CoV-2, and is in contrast with the immunodominant responses directed towards the non-conserved HLA-A^*^02:01-restricted TTD and HLA-A^*^02:01-restricted KLW epitopes report by others [24, 25]. We have previously shown that the efficiency of endogenous antigen recognition can have an impact upon the efficiency of T-cell priming and the immunodominance hierarchy in a number of settings of human viral infection [26, 33, 34]. Proteins critical to intracellular viral function, such as the Epstein Barr Virus Nuclear Antigen 1, often generate subdominant immunity and are not efficiently processed for presentation to CD8^+^ T cells [33, 35]. While limitations in the availability of HLA-matched target cells did not allow us to perform endogenous antigen presentation assays with SARS-CoV-2 infected cells, our observations on antigen abundance from proteomic studies of SARS-CoV-2 infection do suggest that limited antigen abundance could impact T-cell priming against the conserved regions of ORF1ab. It is plausible that this may offer all coronaviruses an immune evasion strategy that limits recognition of highly conserved proteins that are critical for viral replication. More detailed analysis of antigen/epitope abundance and efficiency of T-cell recognition are required to determine this.

One limitation of our study is the relatively homogenous population of COVID-19 convalescent volunteers used to assess T-cell responses. Limitations in the HLA diversity of our cohort likely contributed to the small number of epitopes we discovered. Observations in more genetically diverse individuals may reveal more epitopes. However, considering the devastating impact COVID-19 has had around the world, it seems unlikely that pre-existing cross-reactive CD8^+^ T cells induced by exposure to circulating coronaviruses are having a significant role in protection against severe disease, except in a small proportion of the population. Our observation suggests that the poor immunogenicity of critical regions of ORF1ab limits the efficiency of protection by pre-existing immunity.

## Materials and Methods

### Study participants

This study was performed according to the principles of the Declaration of Helsinki. Ethics approval to undertake the research was obtained from QIMR Berghofer Medical Research Institute Human Research Ethics Committee. Informed consent was obtained from all participants. The inclusion criteria for the study were that participants were over the age of 18, had been clinically diagnosed by PCR with SARS-CoV-2 infection, and had subsequently been released from isolation following resolution of symptomatic infection. A total of 60 participants were recruited in May and June 2020 from the south-east region of Queensland, Australia. Participants ranged in age from 20 to 75, 24 were male and 36 were female, and were a median of 70 (46 – 131) days post-initial diagnosis. Blood samples were collected from all participants to isolate peripheral blood mononuclear cells (PBMC) were isolated. A cohort of SARS-CoV-2 unexposed (n=16) adults was also recruited, to serve as a comparison group.

### Peptides

Overlapping peptide pools (OPP) of nsp12, S, N, and ORF3a were sourced from JPT technologies, Berlin, Germany. Peptides from ORF1ab-C were sourced from Genscript technologies, Hong Kong. The OPPs were 15 amino acids in length overlapped by 11 amino acids sequences to assess the CD8+ T-cell responses. Individual peptides were identified using a two-dimensional matrix-based approach. To carry out epitope fine mapping for the T-cell epitopes, the peptides were trimmed from both N and C terminus. A standard ICS assay was performed to assess all peptide responses.

### In-silico identification of T-cell epitope determinants

The protein sequence of ORF1ab polyprotein (NCBI accession number: QXP49545.1) were retrieved using NCBI (https://www.ncbi.nlm.nih.gov) databases and saved in FASTA format for further analysis. Allele Frequency Net Database (http://www.allelefrequencies.net) was used as a platform to screen allele frequencies across various worldwide populations. Using USA NMDP population we selected n=18 HLA class I (Locus: A, B, C) allele subsets. The ORF1ab polyprotein-HLA class I alleles were used in the prediction of T-cell epitope determinants using Net-MHCpan 4.0 online server (http://www.cbs.dtu.dk/services/NetMHCpan-4.0).

Strong binding affinity peptides of length (8 mer to 13 mer) were selected to design the orf1ab-C peptide pool.

### In-vitro expansion of SARS-CoV-2 specific T cells

PBMC were harvested from peripheral blood within 24 hours of collection. PBMC were incubated with SARS-CoV-2 OPPs (ORF1ab-C, ORF3a, N, S, nsp12) at a concentration of 1μg / mL and incubated at 37°C and 6.5% CO_2_ for an hour. Later, the cells were washed and cultured in 48-well plates for 2 weeks supplemented with recombinant interleukin-2 (IL-2, Charles River Labs, USA) at 20 IU / mL every 2–3 days thereafter. On day 14, T cells were phenotyped and enumerated using the multitest 6-Colour TBNK Reagent (BD Biosciences) and assessed for antigen-specificity using an intracellular cytokine assay. The remaining cells were cryopreserved in culture media with 10% dimethyl sulfoxide (Sigma-Aldrich) and transferred to liquid nitrogen.

### Intracellular cytokine assay

Cultured T cells were stimulated with the cognate SARS-CoV-2 OPPs (ORF1ab-C, ORF3a, N, S, nsp12) and incubated for 4 hours in the presence of Golgiplug (BD Biosciences). Following stimulation, cells were washed and stained with anti-CD8-PerCP-Cy5.5 (eBioscience) and anti-CD4-PE-Cy7 or anti-CD4-Pacific Blue (BD Biosciences) for 30 minutes before being fixed and permeabilised with BD Cytofix/Cytoperm solution (BD Biosciences). After 20 minutes of fixation, cells were washed in BD Perm/Wash buffer (BD Biosciences) and stained with anti-IFN-γ -Alexa Fluor700 or IFN-γ- PE (BD Biosciences) for a further 30 minutes. Finally, cells were washed again and acquired using a BD LSRFortessa with FACSDiva software. Post-acquisition analysis was performed using FlowJo software (TreeStar; FlowJo LLC, Ashland, Oregon). Cytokine detection levels identified in the no-peptide control condition were subtracted from the corresponding test conditions to account for non-specific, spontaneous cytokine production.

### Conservation analysis of peptides derived from pandemic and seasonal coronaviruses

The sequence from SARS-CoV-2 antigen (ORF1ab, N, S, ORF3a) were aligned to alpha coronaviruses (NL63 & 229E) and beta coronaviruses (OC43 & HKU-1) using Clustal omega (https://www.ebi.ac.uk/Tools/msa/clustalo). The T-cell epitopes from SARS-CoV-2 and other seasonal coronaviruses were selected to access cross-recognition and pre-existing immunity.

### Conservation of ORF1ab SARS-CoV-2 isolates by variant

ORF1ab protein sequences for the SARS-CoV-2 (taxid ID 2697049, 419 amino acids) coronaviruses of the following Pango lineages: B1.1.7 (n=9505 sequences from 22.06.2021 until 16.07.2021, reported within as Alpha), B.1.351 (all available n=358 sequences, reported within as Beta), P.1 (all available n=6214 sequences, reported within as Gamma), B.1.617.2 (all available n=327 sequences, reported within as Delta), B.1.427 + B.1.429 (all available n=9850 sequences, reported within as Epsilon), P.2 (all available n=446 sequences, reported within as Zeta), B.1.525 (all available n=487 sequences, reported within as Eta), P.3 (all available n=5 sequences, reported within as Theta), B.1.526 (most recent n=9797 sequences from 07.05.2021 until 16.07.2021, reported within as Iota) and B.1.617.1 (all available n=116 sequences, reported within as Kappa) were obtained from the NCBI virus database (http://www.ncbi.nlm.nih,gov/labs/virus) on the 16^th^ of July 2021. Sequences were aligned using (http://www.fludb.org/brc/home.spg?decorator=influenza). All sequences were used for the alignment. At times, several sequences were unable to be aligned with the reference peptide, and peptides aligned but containing an unknown amino acid (denoted as a X) were excluded from the analysis.

### Cross reactivity and functional avidity of ORF1ab-C T-cell epitopes

T-cell cross-recognition were carried out using ORF1ab-C peptides from SARS-CoV-2, OC43, HKU-1, NL63, 229E. PBMC were grown with SARS-CoV-2 orf1ab-C peptides, cultured for 14 days in the presence of IL-2. On day 14, the ORF1ab-C peptides from OC43, HKU-1, NL63, 229E were used to recall the SARS-CoV-2 peptide to access the cross-recognition by IFN-γ ICS. To assess the functional avidity of ORF1ab-specific T cells, we exposed T cells to ten-fold serial dilution at different concentration (1 μg/mL to 0.00001 μg/ mL) of peptide and performed standard ICS.

### Proteomics analysis

Mass spectrometry raw files were downloaded from PRIDE (PXD018594, PXD017710) and MassIVE repository (MSV000085734) and searched against the UniProt Human protein database and RefSeq SARS-CoV-2 protein database using Sequest HT through Proteome Discoverer (Version 2.2) (Thermo Scientific, Bremen, Germany). Precursor and fragment mass tolerance were set to 10 ppm and 0.02 Da, respectively. Carbamidomethylation of cysteine was set as fixed modifications, while oxidation of methionine was set as a dynamic modification. A false discovery rate (FDR) threshold of 1% was used to filter peptide spectrum matches (PSMs) and peptides. Normalized spectral abundance factor (NSAF) was calculated for all proteins identified. It is calculated as the number of spectral counts (SpC) assigned to a protein by the protein’s length (L), divided by the sum of SpC/L for all proteins in the experiment.

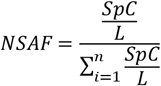

### SARS-CoV-2 microneutralization assay

The protocol for the microneutrlization assay has been previously described [14]. The SARS-CoV-2 isolate hCoV-19/Australia/QLD02/2020 (QLD02) was kindly provided by Queensland Health Forensic & Scientific Services, Queensland Department of Health.

### Statistical analysis

GraphPad Prism 8.2.1 (San Diego, CA) was used to perform statistical analysis. Statistical comparisons between participant groups (unexposed and recovered) were made using unpaired Mann-Whitney U/ Wilcoxon rank-sum tests. Correlative analysis was performed using Pearson correlation co-efficient. Box plots were used to represent median (horizontal line), 25th and 75th percentiles (boxes) and minimum and maximum values (whiskers). *P*<0.05 was considered statistically significant.

## Acknowledgement

The authors would like to thanks Queensland Health Forensic & Scientific Services, Queensland Department of Health who provided the SARS-CoV-2 isolate QLD02. The authors would also like to thanks all the participants who took place in our study. The authors would also like to thank Vivek Venkatram for helping us with Bioinformatics programming. This work was supported by generous donations to the QIMR Berghofer COVID-19 appeal, and the Medical Research Future Fund (MRFF, APP2005654). SS is supported by Australian Government Research Training Program Scholarship and is supported by QIMR-Berghofer Top-Up Scholarship, EJG was supported by an NHMRC CJ Martin Fellowship (#1110429) and is supported by an Australian Research Council DECRA (DE210101479), SVM is supported by QIMR-Berghofer and The University of Queensland scholarship, KRS is supported by an Australian Research Council DECRA (DE180100512), SG is supported by and NHMRC SRF (#1159272).

## Author contribution

SS, KEL, GRA, KRS, MEN, HG, SG RK and CS contributed to the design of the study. SS, KEL, GRA, PC, EJG, SVM, JR, AP, LLT, ZWMT and KYC performed experiments and/or analysis. MAN was responsible for ethics approval and participant recruitment. All authors contributed to drafting of the manuscript.

